# A lipid droplet triacylglycerol lipase governs neutral lipids mobilization and differentiation of *Trypanosoma brucei*

**DOI:** 10.64898/2026.05.09.723952

**Authors:** Perrine Hervé, Yoshiki Yamaryo-Botté, Eloïse Bertiaux, Kathyanna Arnould, Stéphane Claverol, Marc Biran, Emmanuel Tetaud, Frédéric Bringaud, Cyrille Y. Botté, Loïc Rivière

## Abstract

Lipid droplets (LD) are dynamic organelles largely distributed in all living cells that are essential notably for membrane biogenesis and cell proliferation. In *Trypanosoma brucei*, a protozoan parasite responsible for major lethal infectious diseases in humans and cattle, lipid acquisition and mobilization are essential for its survival and adaptation to its different hosts, but the roles of LD in these processes remain largely unknown. To address this gap, we identified and characterized a unique functional orthologue of the human adipose triacylglycerol lipase (ATGL), responsible for the initiation of neutral lipid mobilization, in *T. brucei*. In this study, we present TbPatatin-like (TbPat), the first ATGL ortholog in protozoan parasites. By combining reverse genetics, microscopy and –omics analysis, we deciphered the role of TbPat in lipolysis and neutral lipid maintenance in both insect and mammalian stages of African trypanosomes and imaged its distribution on the LD surface at an unprecedented resolution by using Cryo-Expansion microscopy. We found that inactivation of TbPat in mammalian form *T. brucei* led to a global upregulation of the mitochondrial metabolic proteome; furthermore, these cells appeared remarkably unable to differentiate into the procyclic insect stage. Through its expression and localization, TbPat appears to be at an essential checkpoint linking lipid environment and metabolism to more complex cell programs involving proteome, compartment remodeling and lifecycle completion.

## Introduction

Lipid droplets (LD) are highly dynamic, ubiquitous organelles at the center of cellular lipid flux and metabolism. They possess an atypical architecture consisting of a core filled with neutral lipids, majorly cholesteryl esters (CE) and triacylglycerols (TAG), and bound by a phospholipid monolayer decorated by proteins associated with LD dynamics, lipid metabolism and transport^1^. Originating from the endoplasmic reticulum (ER), the primary function of LD is the storage and distribution of lipids by dynamic associations with other compartments to fuel and regulate cellular demands. Furthermore, due to the toxic potential of lipids, LD are also responsible for the sequestration of excess fatty acids (FA) to prevent lipotoxicity and oxidative stress^2^. For more than a century after their discovery, the role of this compartment has been underestimated, thought as passive lipid inclusions. However, LD have recently emerged as a critical organelle with key implications in metabolic and degenerative diseases, cancers, cellular stress response and host-pathogen interactions^2–5^. Neutral lipids are synthesized in the ER, where they accumulate in oil lenses that form the LD core. Facilitated by several proteins, LD later bud towards the cytosol where they continue to expand and mature by continuous exchanges with the ER^2^. Stored neutral lipids can then be used to provide FA for β-oxidation and precursors for membrane phospholipids through lipolysis and lipophagy, processes mediated by lipases^6^. Lipophagy is the engulfment of a portion or a whole LD to lysosomes, where a broad-substrate lysosomal acid lipase hydrolyses the LD neutral lipid content. Lipolysis is the sequential degradation of TAG by three lipases. The first and rate-limiting step of the reaction is mediated by adipose triglyceride lipase (ATGL), also known as PNPLA2, a TAG lipase from the patatin-like phospholipase family. ATGL localizes to the surface of LD where it converts TAG in diacylglycerol (DAG), releasing free FA^7^. Other lipases then hydrolyze DAG into MAG (HSL) and MAG into glycerol (MGL), releasing FA each time, which then enter different pathways to fuel the cellular energy metabolism or enter membranes as phospholipids^1,6^.

How LD emerge from the ER, interact with other compartments and the regulation of protein targeting and expression to coordinate the different dynamics in LD biology are aspects that are not completely understood. The knowledge on the functions of LD is even more scarce in eukaryotic pathogens such as protozoan parasites, although it represents an emerging interest. Recent studies in other major parasitic protists have demonstrated that LD from both parasites and their mammalian hosts are critical for survival and differentiation in *Plasmodium*^8–10^ and *Toxoplasma*^11–13^, which are responsible for malaria and toxoplasmosis, respectively, by channeling host scavenged FA and timely mobilizing these FA stores to fuel membrane biogenesis and parasite division. Moreover, LD from *Leishmania* spp. and *Trypanosoma cruzi*, which are responsible for leishmaniasis and Chagas disease, respectively, are involved in the production of inflammation factors such as prostaglandin^14–16^. However, the roles of LD in lipid and energy metabolism and the proteins orchestrating the life cycle of LD in these organisms are largely unknown.

*Trypanosoma brucei* is the causative agent of sleeping sickness or African trypanosomiasis in humans and the debilitating disease Nagana in livestock and wild animals. This parasite exhibits a complex life cycle, where it alternates between the bloodstream and other tissues, (brain, adipose tissue, skin, etc.) of mammalians (bloodstream form, BSF) and the midgut (procyclic form, PCF) and salivary glands of its vector, the bloodsucking tsetse fly^17^. To complete this life cycle and survive within its different hosts, *T. brucei* shows unique capacities of adaptation with efficient remodeling of its morphology, proteome and metabolism^18^. In this context, synthesis, acquisition and mobilization of nutrients, including lipids such as FA, is essential to the parasite’s development and survival^19,20^. *T. brucei* can use FA as signaling molecules or as building blocks for complex lipids to maintain and remodel its membrane as well as the lipid anchor that attaches glycosylated proteins called VSG, which form a coat that allows BSF parasites to evade the mammalian host immune response in a process named antigenic variation^20,21^. Besides being a major human and animal pathogen, *T. brucei* is a cellular model from which several ground-breaking discoveries have been made^22^.

To satisfy its lipid requirements, *T. brucei* utilizes different systems depending on its life stage: mammalian form trypanosomes mostly rely on the scavenging and degradation of host lipoproteins^21^ while insect forms use mitochondrial acetate as a precursor to synthesize FA^23,24^ with two essential pathways: a mitochondrial FA synthase (FASII) pathway and an ER FA elongase (ELO) machinery^21,25^. However, despite being at the core of lipid metabolism and trafficking inside organisms, the biological functions of LD, the processes regulating the organelle’s dynamics and interactions, as well as the associated proteins remain mostly uncharacterized in *T. brucei*^26,27^.

In this study, we present the first human ATGL functional orthologue in protozoan parasites. *T. brucei* Patatin-like phospholipase (TbPat) is an active TriAcylGlycerol (TAG) lipase that localizes to the surface of LD through its C-terminal hydrophobic domain. We used the combination of ultrastructure expansion microscopy (U-ExM) with cryo-fixation (Cryo-ExM) and showed for the first time the preservation of LD during expansion, allowing visualization of LD-associated proteins and LD-organelle contact sites with accessible super-resolution microscopy. The turnover of TbPat is tightly regulated by parasites and its stability significantly increases after oleate-induced LD formation. Overexpression of TbPat leads to a significant decrease of TAG levels and LD number in *Trypanosoma* parasites, while inactivation of the enzyme results in enlarged LD and a dysregulation of the balance of neutral lipids in bloodstream and procyclic parasites, including a significant increase of TAG levels. We identified the canonic lipase active serine as responsible for TbPat activity, and expression of a catalytic mutant version of the protein is severely deleterious for cells in presence of oleate. Inactivation of TbPat also leads the upregulation of several mitochondrial metabolic pathways of dividing mammalian form parasites, concurrent with an inability to differentiate into the procyclic stage.

These findings identify TbPat as a key regulator of LD dynamics and neutral lipids balance in trypanosome parasites, provide new insights on the roles of LD in maintaining energy homeostasis and bring to light a new, unexplored function of the organelle as a regulator of parasite differentiation. Finally, they demonstrate the utilization of trypanosomes as a new, effective model to study lipid droplet biology and dynamics.

## Results

### Trypanosomes encode for a putative Patatin-like phospholipase ortholog to Human Adipose Triglyceride Lipase

The characteristics of the Patatin fold (PFAM 01734) has been previously described, particularly in the plant lipase PAT17 for which the crystal structure was resolved^28,29^. This domain displays an α/β class protein fold with three α/β/α layers in which a central six-stranded β-sheet is sandwiched between α-helices front and back. The central β-sheet contains five parallel strands and an antiparallel strand at the edge of the sheet. Another important feature of patatin-like proteins is the lipase motif, characterized by a Ser-Asp catalytic dyad. A combined text search in the African trypanosome *Trypanosoma brucei* for ‘Patatin’ and motif pattern search for ‘G.X.S.X.G’ (lipase motif) in the TriTryp database (https://tritrypdb.org), yielded a unique candidate with the identifier Tb927.4.1780, thus called TbPat (Fig. 1a). According to comparative genome analysis, this gene/protein is syntenic in kinetoplastids, exhibiting a high degree of similarity (>60%) between species, among them *Leishmania infantum* and *Trypanosoma cruzi*, two other deadly human pathogens. Protein sequence alignment of TbPat with PAT17 revealed a low level of similarity, except for the lipase motif identified by a serine residue at position 68 and an aspartate residue at position 194 (Extended Data Fig. 1). Attempts to express TbPat in a heterologous system were unsuccessful, thus, to gain structural insights, we employed AlphaFold3 to model it. As shown, the model reproduces with high confidence the several sheets and helix and the two catalytic residues in a structural close proximity as well as the two portals necessary for substrate accessibility and subsequent enzymatic activity (Fig. 1b). This suggests that TbPat likely possesses enzymatic activity. α/β Hydrolases containing a Patatin domain are subdivided in different groups based on sequence similarities. To determine which group TbPat belongs to, we reconstructed a phylogenetic tree as previously done^7^. As shown, TbPat is evolutionary close to HsATGL and consequently clusters with the Adipose TriGlyceride Lipase family (Extended Fig. 1). Interestingly both TbPat and HsATGL harbor the same α/β fold with 4 parallel β-sheets followed by one anti-parallel, which is slightly different from the 5 versus 1 sheet observed in PAT17 (Fig. 1c). Furthermore, a putative hydrophobic domain (from amino acid 298 to 321) could be predicted in TbPat. In HsATGL, this domain may favor interactions with lipids^30^. We thus inferred from all analysis, predictions and observations that the protozoan TbPat protein is a HsATGL functional orthologue.

**Fig. 1:**
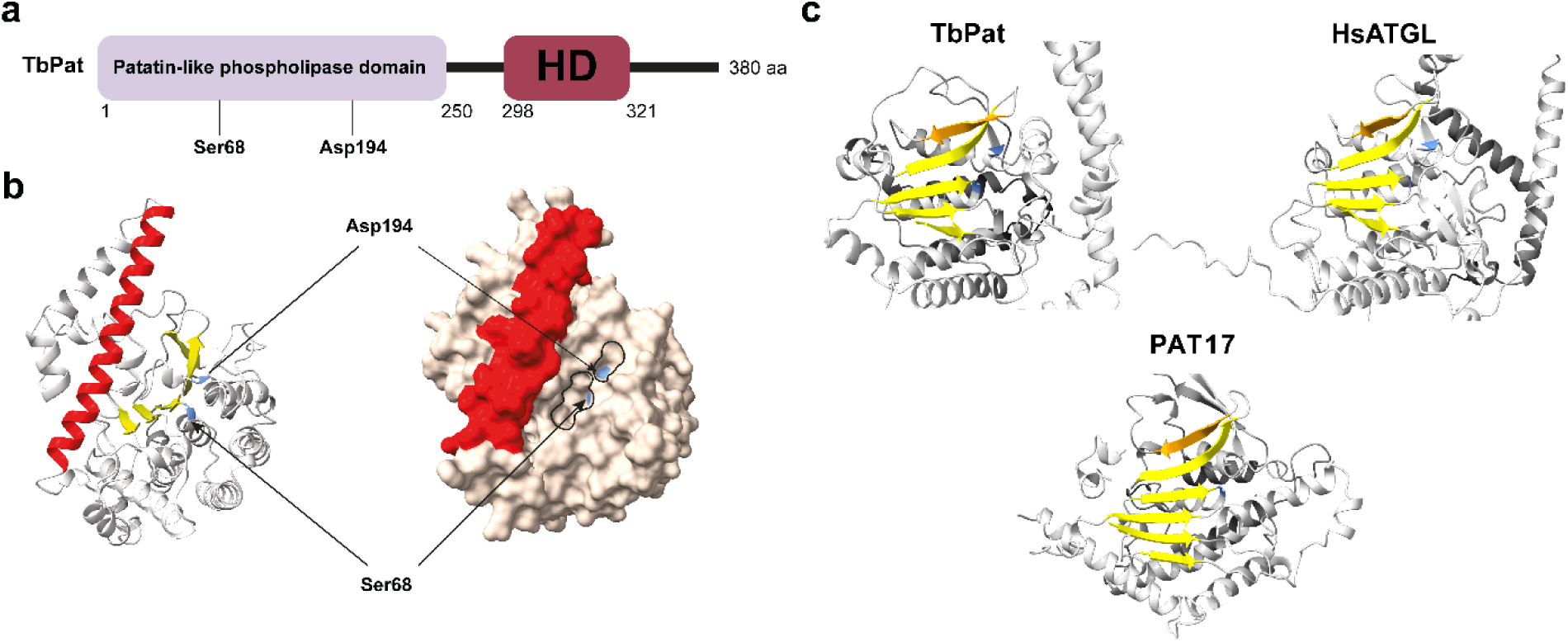
TbPat, a putative Patatin-like phospholipase. **a**, Scheme depicting the domains of TbPat. Purple box: patatin-like domain with critical catalytic residues Ser 68 and Asp 194. Red box: Hydrophobic Domain (HD). **b**, Alphafold3 model of TbPat visualized using ChimeraX 1.9. Left model, Structure showing helix and beta sheets organization. HD corresponds to the red helix, alpha/beta hydrolase fold is in yellow, Ser68 and Asp194 are in blue; Right model, volume representation showing the pocket with the catatylic residues rounded in black. **c**, Predicted structures emphasizing the alpha/beta fold domains of PAT17 (*T. solanum*), ATGL (*H. sapiens*) and TbPat. PAT17 has 5 parallels and one anti parallel beta sheet arrangement, both ATGL and TbPAT have 4 parallels and one anti-parallel beta sheet.

### TbPat localizes to lipid droplets through its C-terminal hydrophobic domain

Tryptag, a published genome-wide database of the subcellular localization of *T. brucei* proteins, allowed us to observe that N– or C-terminal mNeonGreen-tagged versions of TbPat may localize to LD^31^. As the inferred localization is made solely by visual deduction, we first intended to verify this annotation. Thus, we conditionally expressed a GFP-tagged version of the protein (GFP-TbPat) in procyclic trypanosomes (Fig. 2a); expression of the fusion protein after tetracycline (tet) induction was confirmed by Western-blot with the anti-GFP (Fig. 2b). Immunofluorescence of tet-induced cells showed that GFP-TbPat consistently co-localized with the LD marker Lipi-blue, confirming TrypTag localization (Fig. 2d).

**Fig. 2:**
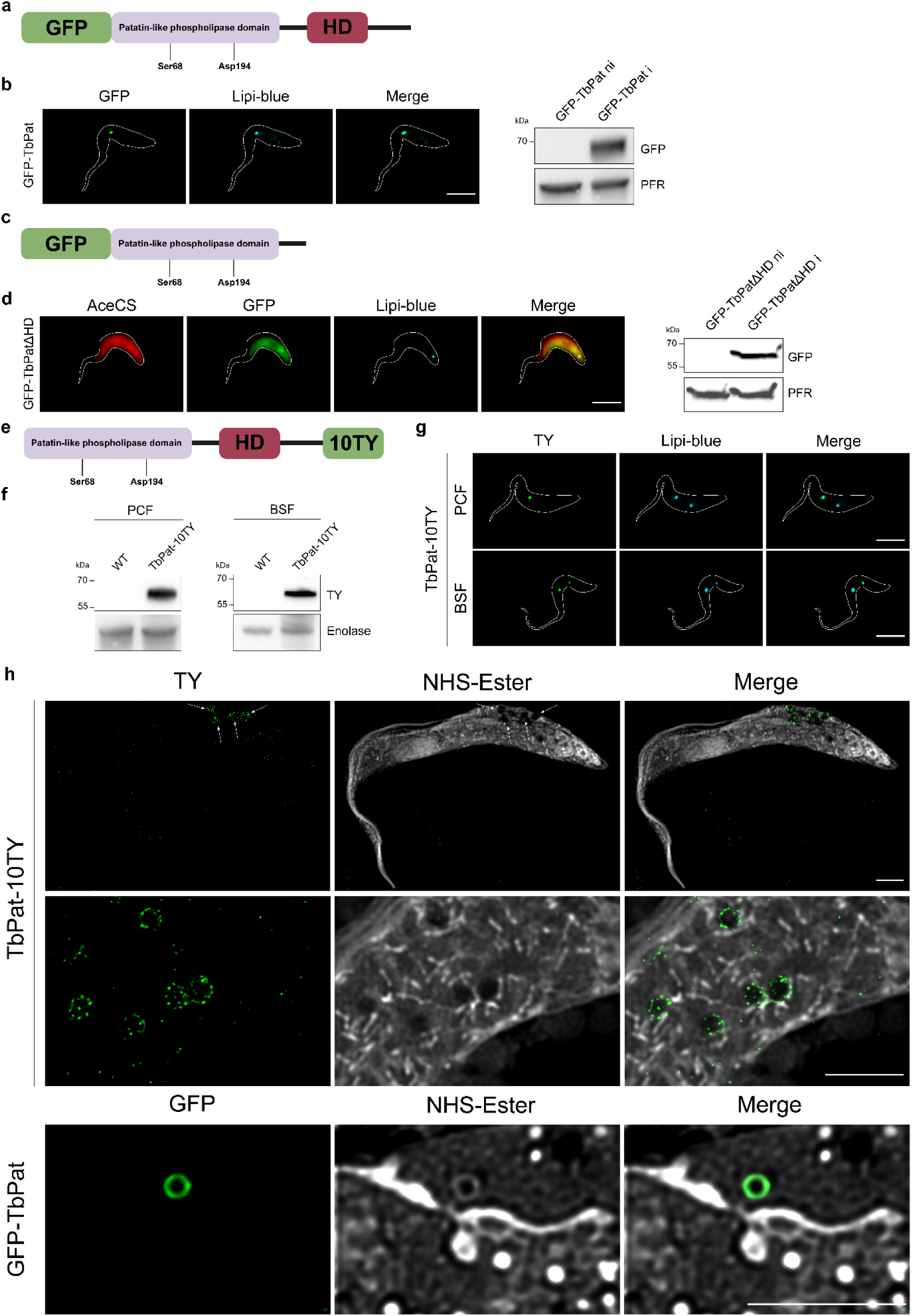
TbPat is a lipid droplet-bound protein. **a**, A schematic representation of TbPat fused at the N-terminus with a GFP tag in the plew100 vector. **b**, Fluorescence image of tetracycline-induced fixed GFP-TbPat. The lipid droplet dye Lipi-blue was used to specifically stain lipid droplets (left) and Western-blot analysis of non-induced and induced GFP-TbPat (right). Protein loading was controlled using the trypanosomal paraflagellar rod (PFR) protein. **c,** A schematic representation of TbPat lacking the C-terminal containing the hydrophobic domain (TbPatΔHD). The sequence was fused to GFP and inserted in the plew100 vector. **d**, Immunofluorescence of tetracycline-induced GFP-TbPatΔHD with cytosolic *T. brucei* Acetyl-CoA synthetase (AceCS) and Lipi-blue (left) and Western-blot analysis of non-induced and induced GFP-TbPatdHD (right). Protein loading was controlled using PFR. **e**, A schematic representation of endogenous TbPat fused at the C-terminus with a 10TY tag in both alleles. **f**, Western-Blot of wild-type and TbPat-10TY procyclic form (left) and bloodstream form (right) parasites. Protein loading was controlled using Enolase. **g**, Immunofluorescence of procyclic (top) and bloodstream (bottom) TbPat-10TY parasites with Lipi-blue. **h**, Cryo-Expansion microscopy of TbPat-10TY (top, middle) and GFP-TbPat (bottom) cells. NHS-Ester labels free amino groups of proteins. PCF, procyclic form; BSF, bloodstream form. Scale bars: 5 µm.

In contrast to other compartments like peroxisomes or mitochondria, there is no canonical targeting sequence for protein addressing to LD. Nevertheless, it has been demonstrated in higher eukaryotic organisms that the presence of hydrophobic sequences is essential for correct LD targeting for several proteins^32^. Consistent with those observations, previous analysis of ATGL showed that removal of its hydrophobic domain (ATGLΔHD) prevented association with the LD single-layer membrane^30^. To investigate this question in TbPat, we conditionally expressed a truncated version lacking the predicted hydrophobic domain of the GFP-tagged TbPat (GFP-TbPatΔHD) (Fig. 2c). Protein expression was controlled by Western-blot ting and imaged by immunofluorescence with the anti-GFP (Fig. 2d). As previously observed with ATGL, tet-induced GFP-TbPatΔHD no longer localized to the LD surface but was instead diffused in the cytosol and colocalized with the cytosolic protein AcetylCoA Synthetase (AceCS)^23^.

It cannot be ruled out that the presence of the GFP/mNeon-Green or the inducible character of the GFP-TbPat fusion protein may interfere with proper addressing of TbPat, leading to a mislocalization. We thus engineered, by using a marker-free CRISPR/CAS9 approach recently optimized by our group^33^, a new cell-line by inserting another tag (10TY^34^) directly at the 3’ extremity of both alleles of the endogenous *TbPat* genomic locus in procyclic and mammalian form parasites (Fig. 2e). The correct integration of the 10TY cassette was checked by PCR amplification, and Western-blot analysis with the anti-TY confirmed the expression of the 10TY-tagged TbPat (TbPat-10TY) at the expected molecular weight (56 kDa) (Fig. 2f). Immunofluorescence analysis of TbPat-10TY parasites with the anti-TY still revealed a clear association of the protein with Lipi-blue in both lifecycle stages (Fig. 2g). Overall, these experiments show that N– or C-terminal-tagged TbPat localizes to *T. brucei* lipid droplets. Moreover, interaction with the LD surface may be attributed to the presence of an C-terminal hydrophobic domain, as previously observed for HsATGL^30^. It is important to note that it is only the second characterized protein with such a localization in trypanosomes, after the kinase TbLDK^26^.

### Super-resolution visualization of TbPat by Cryo-Expansion Microscopy reveals discrete localization at the LD surface

To get a better resolution of TbPat localization on the LD surface, we employed Ultrastructure Expansion Microscopy (U-ExM)^35^, a super-resolution technique that has already been employed in *T. brucei* to locate proteins at a nanoscale resolution^36,37^. However, “conventional” U-ExM does not support the study of LD proteins as these structures are not maintained during the expansion process (Extended Data Fig. 2a). Thus, we combined this approach with cryo-freezing (Cryo-ExM), a method that preserves the cell architecture and limits artifacts compared to chemical fixation^38^. In this approach, like in conventional U-ExM, neutral lipids are removed and in consequence Lipi-blue, Nile Red or BODIPY dyes are not effective in staining LD post-expansion. To bypass this technical limitation, we used NHS-Ester, which covalently labels the amino terminus of proteins, to allow us to infer the position of LD in expanded cells (Fig. 2h). Cryo-ExM with the anti-GFP or anti-TY on conditionally expressed GFP-TbPat and endogenous TbPat-10TY cell lines, respectively, showed that endogenous TbPat-10Y is not uniformly distributed on the LD surface (Fig. 2h, top and middle panels), whereas conditionally expressed GFP-TbPat is (Fig. 2j, lower panel). LD are formed from the ER and remain closely associated with the compartment after biogenesis, but they are also known to form contact with other organelles such as mitochondria and peroxisomes^39^. ER-resident protein BiP staining in *T*.

*brucei* showed that the architecture of the compartment was preserved during expansion. Interestingly, TY-tagged TbPat is observed on the border of the ER but does not overlap with the compartment, suggesting that the two organelles remain in close contact after LD budding but that TbPat is physically separated from the ER (Extended Data Fig. 2b). We also found that LD were closely associated with the unique mitochondrion of *T. brucei*, as well as glycosomes, specialized peroxisomes of kinetoplastids, suggesting previously undiscovered interactions between these compartments in the parasite (Extended Data Fig. 2c).

Hence, we demonstrated that the use of Cryo-ExM allows accessible, super-resolution visualization of LD proteins and revealed that endogenous TbPat does not uniformly decorate the LD surface, a phenotype never observed before in ATGL or other LD-associated lipases.

### TbPat abundance regulates LD number through lipolysis

Previous work from our group showed that under normal conditions *T. brucei* has fewer than 2 LDs per cell, but adding oleate to the culture medium induces LD formation, increasing the number by 4-fold^40^. To understand the role of TbPat in LD dynamics, we investigated the pattern of abundance of the endogenous tagged protein following oleate feeding. After 24 hours of fatty acid supplementation, we observe the formation of new lipid droplets in procyclic and mammalian form parasites (Fig. 3a), coinciding with a significant increase of TbPat abundance both at the mRNA and protein levels (Fig. 3b and 3c). A heterogeneous signal of TbPat on the LD surface could once again be observed, suggesting its specific interaction with lipids and/or other proteins. To date, TbPat is the first protein from trypanosomes known to increase in abundance during exogenous fatty acid feeding, hinting at an unknown regulatory system for LD biology and lipid acquisition in these organisms. To gain further clues on the function of TbPat we investigated the effects, under the same oleate feeding conditions, in the GFP-TbPat cell-line (Fig. 3d). We observed that in contrast to the TbPat-10TY cell line, addition of oleate had no effect on the levels of the tetracycline-controlled protein abundance (Fig. 3e). However, we noticed a significant decrease in the mean number of LD in tet-induced GFP-TbPat parasites compared to non-induced cells, which was more dramatic after oleate feeding (Fig. 3f), while LD area was not affected (Fig. 3g). Furthermore, tet-induction of a supplemental copy of the protein caused a slight growth delay when parasites were maintained in a culture medium supplemented with fatty acids (Fig. 3h). GFP fluorescence of induced cells was evaluated daily during the growth curves (Extended Data Fig. 2e)

**Fig. 3:**
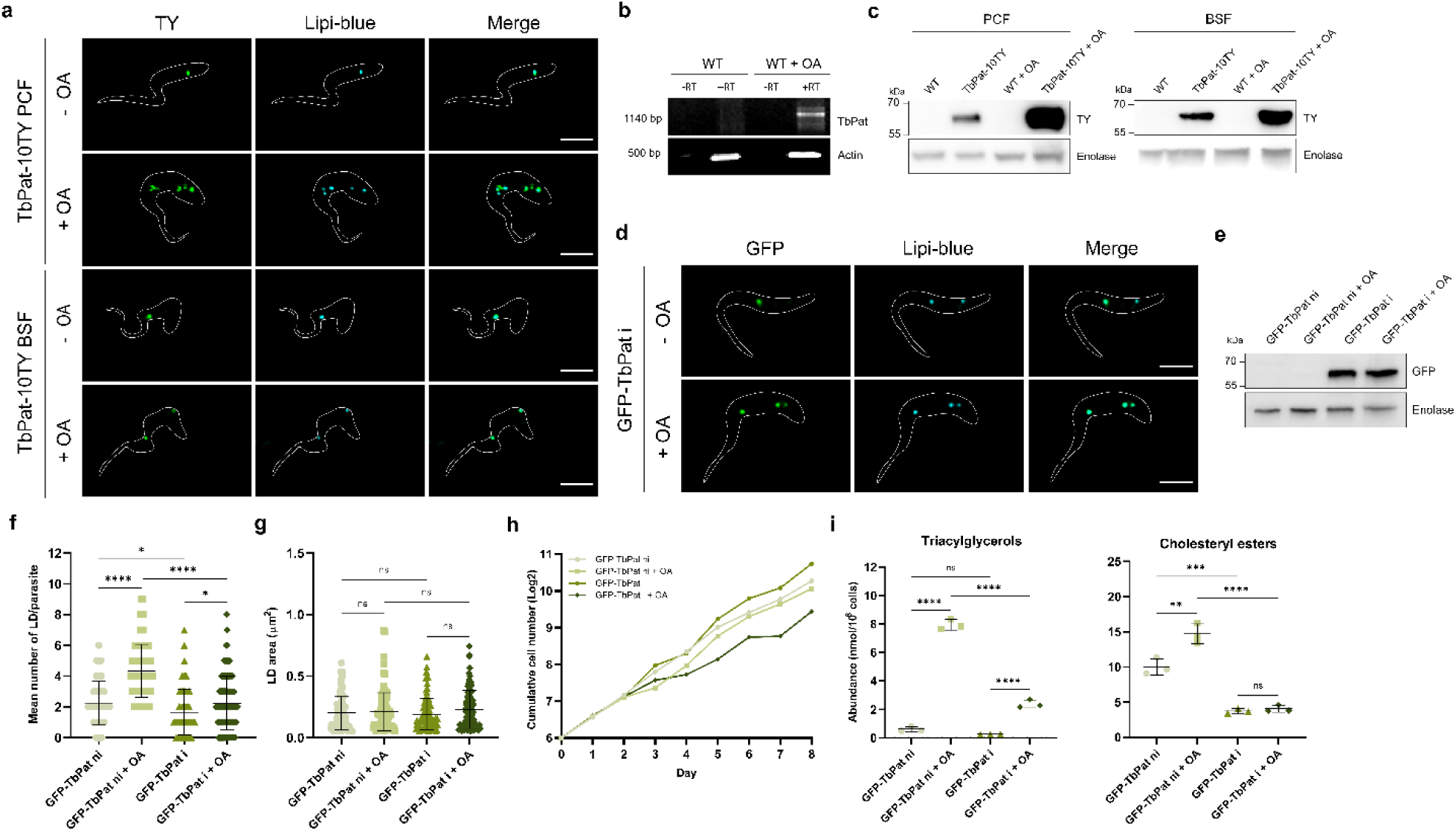
TbPat abundance drives lipolysis through an environment-dependent regulation. **a,** Immunofluorescence of TbPat-10TY procyclic (top) and bloodstream (bottom) parasites untreated or treated with 400 μM oleic acid for 24 hours. **b,** RT-PCR analysis of TbPat in untreated or oleic acid-supplemented procyclic cells. PCR on TbPat and TbActin was performed on purified RNA before (-)and after (+) reverse transcription. Actin was used as loading control. Western-blot analysis of wild-type and TbPat-10TY procyclic (top) and bloodstream (bottom) cells. **c,** Western-blot analysis of wild-type and TbPat-10TY procyclic (top) and bloodstream (bottom) cells. **d,** Fluorescence images of induced untreated or oleic acid-treated fixed GFP-TbPat cells. The expression of the fusion protein was induced for 5 days before addition of 400 μM oleic acid. **e,** Western-Blot analysis of non-induced and induced GFP-TbPat, without or with 400 μM oleic acid supplementation. **f, g** Effect of GFP-TbPat induction and oleate supplementation on the mean number of lipid droplet per parasite (**f**) and lipid droplet area (**g**). n = 100 parasites/condition (**f**) or n = 100 lipid droplets/condition (**g**). **h,** Growth curves of non-induced and induced GFP-TbPat, without or with oleate treatment. Oleic acid was added after every dilution. The curves represent the mean growth of 3 biological replicates **i,** Effect of GFP-TbPat induction and oleate supplementation on triacylglycerols (left) and cholesteryl esters (right) abundances. Abundances were normalized to 10^8^ cells. n = 3 biological replicates. All images were analyzed using ImageJ. The graphical data are presented as means + sd. ns = not significant, *, p-value <0.05; **, p-value <0.005; ****, p-value <0.0001. Statistical significance was determined by one-way ANOVA with Tukey’s multiple comparisons test (95% confidence interval). Graphs were generated using GraphPad Prism v8. 0.2. ni, non-induced; i, induced. OA, oleic acid. Scale bars: 5 µm.

To understand how TbPat impacts LD dynamics, we performed lipidomic analysis on non-induced and tet-induced GFP-TbPat procyclic cells cultivated with or without oleic acid for 24 hours. Since neutral lipids are the main components of LD, we initially focused our analysis on the two main neutral lipid classes present in LDs, TriAcylGlycerols (TAG) and Cholesteryl Esters (CE). In standard culture conditions, the amount of CE in non-induced GFP-TbPat PCF trypanosomes is almost 10-fold higher than that of TAG (Fig. 3i). Added oleate was incorporated into both TAG and CE, but while the overall CE levels remained unchanged, TAG content increased 4-fold (Fig. 3i), indicating that the main pathway for fatty acid internalization in PCF *T. brucei* is through TAG synthesis; diacylglycerol (DAG), monoacylglycerol (MAG) and free fatty acids levels also increased, but to a lesser extent (Extended Data Fig. 3a). Interestingly, the amount of TAG is significantly lower in the tet-induced GFP-TbPat cells when compared to non-induced cells, regardless of oleic acid supplementation (Fig. 3i), and was not accompanied by a significant accumulation of DAG or MAG (Extended Data Fig. 3a). In addition, the increase in CE was also less pronounced after tet-induction of GFP-TbPat expression (Fig. 3i). These observations suggest that the decrease in TAG and CE abundance as well as the decrease of the number of LD per cell, associated with the expression of an additional *TbPat* gene copy, was likely due to an enhanced hydrolysis rather than a reduced TAG synthesis. Therefore, we propose that TbPat is an enzyme that favors lipolysis and whose association with LD is tightly controlled upon lipid environment.

### Loss or catalytic inactivation of TbPat disrupts lipid droplet homeostasis

To explore more deeply the role of TbPat in lipid droplet biology, we inactivated TbPat in PCF parasites by inserting stop codons in both *TbPat* alleles using a markerless CRISPR/Cas9 approach, as recently decribed^33^. Briefly, we inserted a short cassette containing STOP codons in all three open reading frames surrounding the HpaI restriction site inside the coding sequence of TbPat (ΔTbPat) (Fig. 4a). Integration of the cassette in clonal populations was confirmed after PCR amplification of the *TbPat* sequence followed by HpaI digestion. Comparison of the area and number of LD in WT cells and ΔTbPat parasites showed a significant 3-fold increase in LD area in ΔTbPat cells (Fig. 4b and 4e). Furthermore, while WT cells were able to produce new LD after oleate addition to store the excess produced lipids, ΔTbPat parasites displayed an important defect in the generation of new LD (Fig. 4b and 4f). To ensure that this phenotype was only attributed to the loss of TbPat, we generated an add-back cell line by introducing the GFP-TbPat plasmid in ΔTbPat PCF parasites (ΔTbPat::GFP-TbPat). Surprisingly, in this cell line, GFP-TbPat no longer localized to the LD surface, but was instead consistently found in the endoplasmic reticulum (Fig. 4d, Extended Data Fig. 2d). Tet-induction of the fusion protein still led to a marked decrease in LD area (Fig. 4e), but also an important decrease of the mean number of LD per parasite compared to WT cells (0.79 versus 2.15, respectively) (Fig. 4f), as Lipi-blue staining of tet-induced cells showed a unique, diffuse marking of neutral lipids in cells, with very few LD (Fig. 4d). LD-associated proteins are targeted to the LD surface through two different pathways: ERTOLD (Endoplasmic Reticulum TO Lipid Droplet) or CYTOLD (Cytosol TO Lipid Droplet). ER localization of TbPat indicates that the enzyme may use the ERTOLD pathway to access the LD monolayer, which has previously been hinted at as a mechanism for ATGL targeting to LD^41^. This possible ER retention also echoes findings in yeast, where TAG lipases relocalize to the ER in the absence of neutral lipid storage^42^. Western-blot analysis showed that GFP-TbPat was more abundant in oleate-fed cells (Fig. 4c), despite its expression being tetracycline-dependent, suggesting post-translational stabilization of TbPat under lipid-rich conditions.

**Fig. 4:**
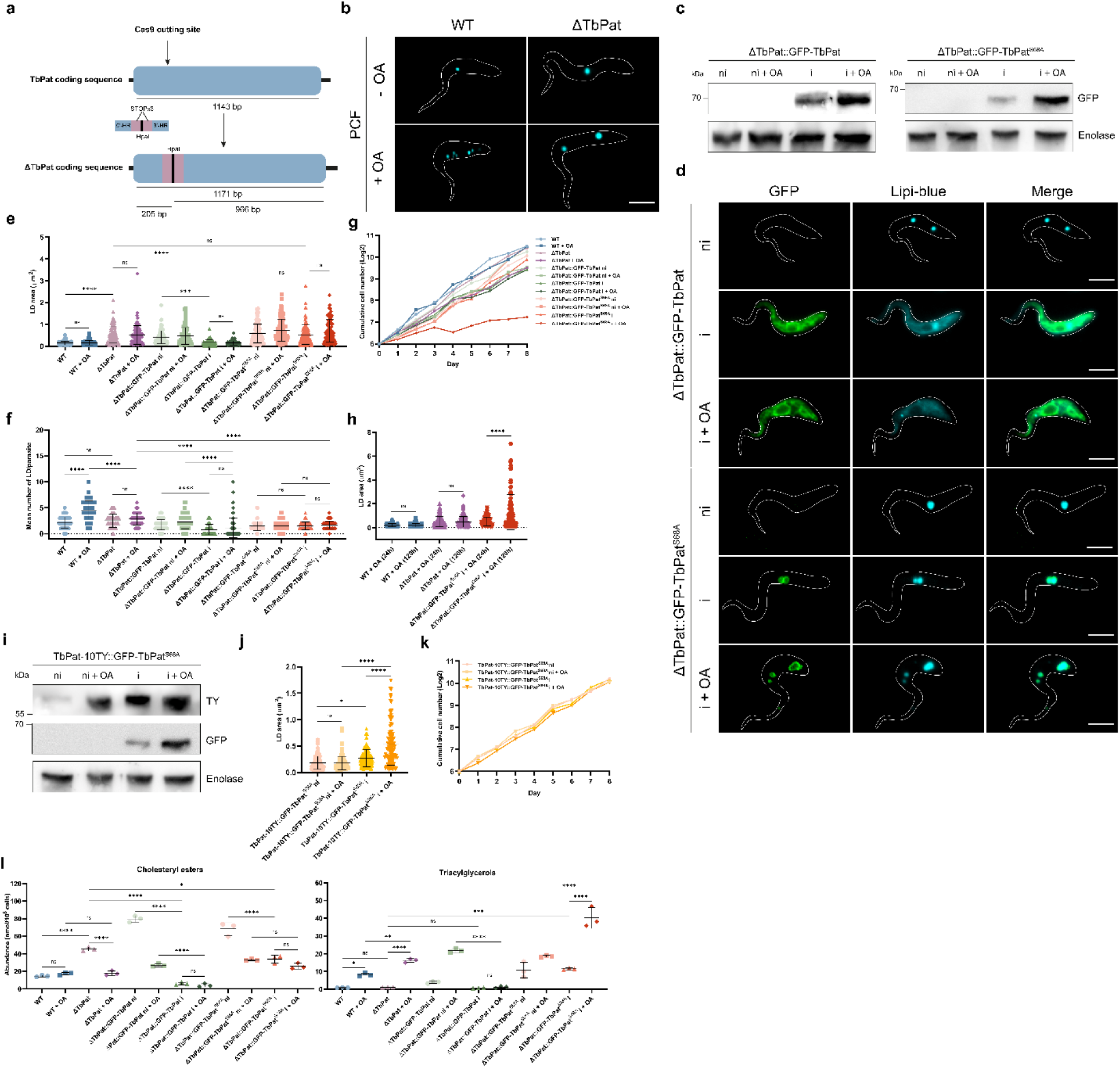
Precise regulation of TbPat expression and catalytic is necessary for *T. brucei* lipid homeostasis and survival. **a**, A schematic representation the TbPat inactivation strategy by insertion of a short cassette containing a succession of STOP codons surrounding the HpaI restriction site flanked by around 40 bp of homology (HR) to the 5 and 3’of TbPat. **b,** Lipid droplet imaging of untreated or oleic acid-treated procyclic wild-type and ΔTbPat parasites using Nile Red. **c,** Western-blot analysis of non-induced or induced ΔTbPat::GFP-TbPat (left) and ΔTbPat::GFP-TbPat^S68A^ (right) with or without oleate treatment. **d,** Fluorescence image of ΔTbPat::GFP-TbPat (top) and ΔTbPat::GFP-TbPat^S68A^ (bottom) non-induced or induced cells. **e,** Quantification of lipid droplet area of untreated or oleic acid-treated wild-type, ΔTbPat, ΔTbPat::GFP-TbPat and ΔTbPat::GFP-TbPat^S68A^ parasites. n = 100 lipid droplets/condition. **f,** Quantification of lipid droplet area of wild-type, ΔTbPat, and ΔTbPat::GFP-TbPat^S68A^ parasites treated with oleic acid for 24 or 120 hours. n = 100 lipid droplets/condition. **g,** Quantification of the mean number of lipid droplet per parasite in wild-type, ΔTbPat, ΔTbPat::GFP-TbPat and ΔTbPat::GFP-TbPat^S68A^ parasites. n = 100 parasites/condition. **h,** Growth curves of wild-type, ΔTbPat, ΔTbPat::GFP-TbPat and ΔTbPat::GFP-TbPat^S68A^ parasites, without or with oleate treatment. Oleic acid was added after each dilution. **i,** Western-blot analysis of non-induced or induced TbPat10TY::GFP-TbPat^S68A^ without or with oleate treatment**. j,** Quantification of lipid droplet area of non-induced or induced TbPat10TY::GFP-TbPat^S68A^ parasites, without or with oleate treatment. n = 100 lipid droplets/condition. **k,** Growth curves non-induced or induced TbPat10TY::GFP-TbPat^S68A^ parasites, without or with oleate treatment. Oleic acid was added after every dilution. **l,** Quantification of cholesteryl esters (left) and triacylglycerols (right) abundances in wild-type, ΔTbPat, ΔTbPat::GFP-TbPat and ΔTbPat::GFP-TbPat^S68A^ parasites. Abundances were normalized to 10^8^ cells. n = 3 biological replicates. **j,** Western-blot analysis of non-induced or induced GFP-TbPat^S68A^ in TbPat-10TY cells with or without oleate treatment. The curves represent the mean growth of 3 biological replicates. All images were analyzed using ImageJ. The graphical data are presented as means + sd. ns = not significant, *, p-value <0.05; **, p-value <0.005; ***, p-value <0.001; ****, p-value <0.0001. Statistical significance was determined by one-way ANOVA with Tukey’s multiple comparisons test (95% confidence interval). Graphs were generated using GraphPad Prism v8.0.2. ni, non-induced; i, induced. OA, oleic acid. Scale bars: 5 µm.

To probe the functional importance of TbPat’s catalytic core, we engineered a new cell line by introducing a serine-to-alanine mutation (S68A) in the Patatin domain of the GFP-TbPat sequence, which was then transfected in ΔTbPat parasites. As opposed to ΔTbPat::GFP-TbPat parasites, LD area and number remained unchanged in the tet-induced catalytic mutant cell line when compared to the parental ΔTbPat strain, indicating that the lipase-canonical catalytic serine is essential for TbPat enzymatic activity (Fig. 4e and 4f). Classical immunofluorescence and Cryo-ExM microscopy revealed that the mutant protein localized broadly around still-enlarged LD and accumulated at LD-LD contact sites (Fig. 4d, Extended Data Fig. 2c). Interestingly, Western-blot analysis with the anti-GFP showed that, similarly to ΔTbPat::GFP-TbPat, protein abundance was more important in oleate-fed cells (Fig. 4c), further reinforcing a hypothetical regulation of the turnover of TbPat depending on the lipid environment, which is independent of its activity. While inactivation of TbPat only had a moderate effect on parasite growth during oleate supplementation, which was still observed after expression of GFP-TbPat in the TbPat null background (Fig. 4g), oleic acid feeding of cells expressing the catalytic mutant had major consequences on parasite growth, concurrent with a gradual LD enlargement over 5 days of oleate feeding, which was not observed in ΔTbPat (Fig. 4g, Fig. 4h). GFP fluorescence of the induced cell lines was controlled daily during the growth curves (Extended Data Fig. 2e).

LD enlargement was also observed when expressing the catalytic mutant in the endogenously-tagged TbPat-10TY cell line, where tet-induction of the fusion protein led to a concomitant increase of endogenous TbPat (Fig. 4i, Fig. 4j), but did not have any effect on cell growth (Fig. 4k), suggesting that endogenous TbPat is positively regulated to compensate for the deleterious effect of the catalytic mutant protein to maintain a minimal flow of TAG hydrolysis.

These results underline the necessity of a tight regulation of TbPat expression by trypanosomes and its activity in lipid modulation for parasite survival.

### TbPat is a TAG lipase responsible for the regulation of the neutral lipid balance of *T. brucei*

Lipidomic analysis on WT and ΔTbPat parasites in PCF cells cultivated with or without oleate for 24 hours was performed to assess how the absence of the protein would impact lipid dynamics in different conditions. Under standard conditions of culture, during oleate feeding of ΔTbPat cells, CE levels rose by 3-fold (Fig. 4l) with no significant changes in the relative abundances of its different molecular species (Extended Data Fig. 3b). While TAG levels were not impacted by the loss of TbPat in these conditions compared to WT cells (Fig. 4i, right), addition of oleic acid in ΔTbPat cells led to a clear inversion of the abundances of CE and TAG: CE levels dramatically decreased to return to WT-like levels while TAG levels increased by 14-fold, significantly higher than in WT oleate-fed parasites (Fig. 4l). This result can be linked to the lack of LD formation observed in ΔTbPat oleate-supplemented cells and shows it is likely due to a replacement of the LD content rather than a defect of organelle biogenesis in these conditions (Fig. 4f). Reintroduction and tet-induction of the TbPat sequence in the null background restored WT-like content of TAG and CE (Fig. 4l). Furthermore, oleate feeding did not result in significant increase of either neutral lipid after tet-induction (Fig. 4l), which explains the low number of LD observed in these parasites (Fig. 4d, Fig. 4f). Expression of the catalytic mutant resulted in a drop of CE levels, hinting at a separate, unknown regulatory mechanism for cholesterol esterase activity (Fig. 4l). However, TAG abundance did not decrease and the neutral lipid even significantly accumulated after oleate feeding (Fig. 4l), related to the LD size increase observed after several days of oleate feeding due to extensive TAG storage (Fig. 4e, Fig. 4h).

Collectively, these results confirm that TbPat is a TAG lipase. Its inactivation causes a general dysregulation of neutral lipids balance and mobilization in trypanosomes and suggest that global regulation of neutral lipid balance by the mean of TAG and CE in *T. brucei* is mainly mediated by TbPat.

### Loss of TbPat induces a remodeling of the mitochondrial proteome and prevents differentiation of bloodstream form *Trypanosoma brucei*

In BSF trypanosomes, like PCF parasites, inactivation of TbPat led to significant increase of LD size and a defect in LD formation during oleate feeding (Fig. 5a, Fig. 5b, Fig. 5c). Contrary to PCF cells, addition of oleic acid in WT cells had no significant impact on either CE or TAG levels, but rather on DAG abundance (Fig. 5d), indicating different regulations of fatty acid incorporation depending on the life stage of the parasite. Loss of TbPat did not impact global CE, DAG or TAG levels, but the relative abundance of several molecular species was inverted compared to WT cells: abundances of FA with shorter carbon chains, from C16 to C20, were significantly reduced in ΔTbPat cells while levels of the very long unsaturated chains of C22 dramatically increased. We observed this inverted pattern for both CE and TAG chains (Extended Data Fig. 3c). By comparing the relative abundance of CE molecular species in BSF and PCF cells, we found that the profile of BSF ΔTbPat parasites was closer to WT PCF cells than WT BSF (Extended Data Fig. 3d). Furthermore, after oleate feeding, the overall abundance of CE in ΔTbPat significantly decreased while, at the same time, TAG levels dramatically increased, as already observed in PCF cells, confirming TbPat TAG lipase activity. Addition of oleic acid was lethal to both WT and ΔTbPat cells after several days of culture (Fig. 5e) which is consistent with a previous report^43^.

**Fig. 5:**
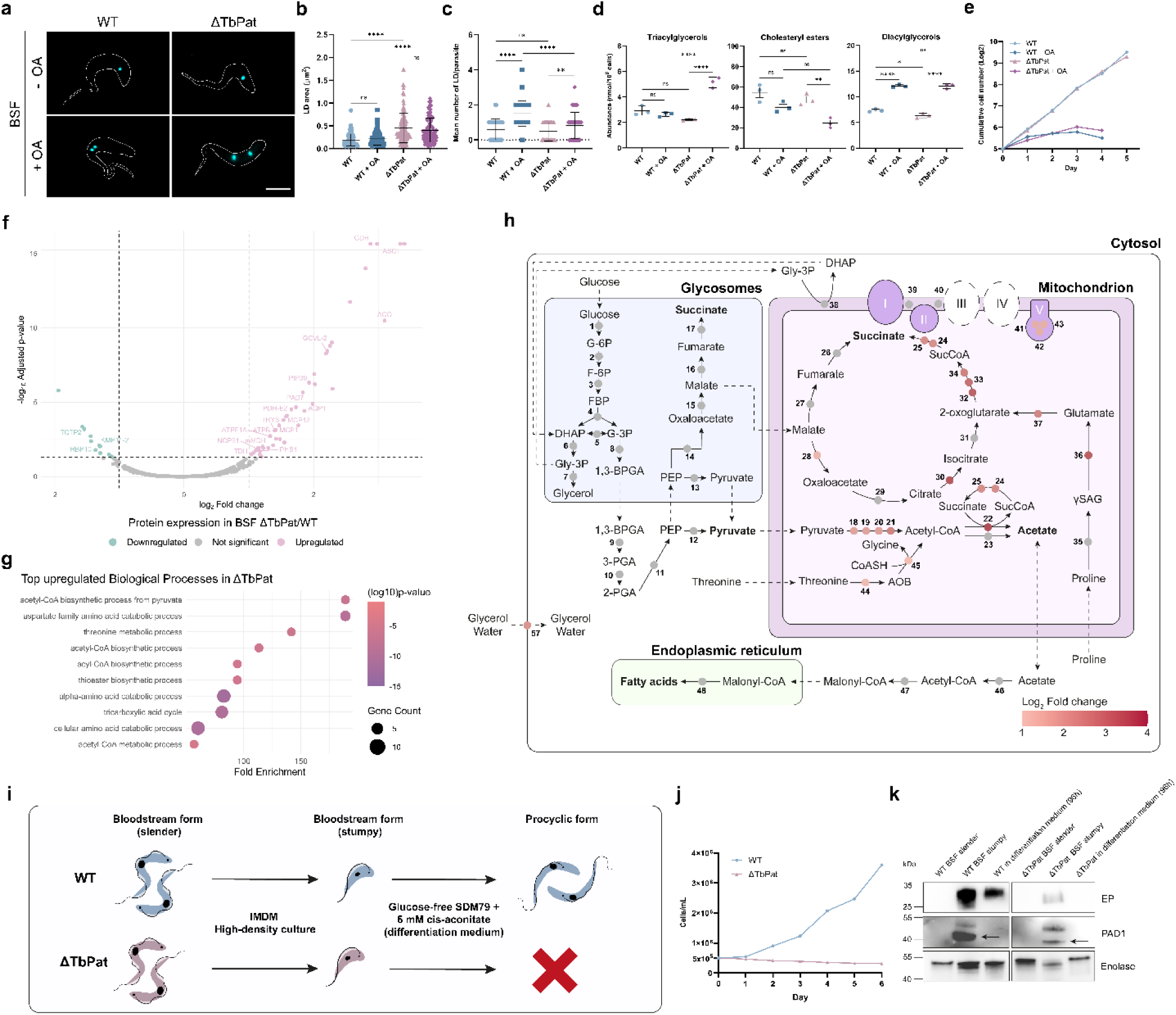
Loss of TbPat causes a remodeling of mammalian form *T. brucei* metabolic proteome and prevents lifecycle progression. **a,** Lipid droplet imaging of untreated or oleic acid-treated procyclic wild-type and ΔTbPat parasites using Nile Red. **b,** Quantification of lipid droplet area of oleic acid-untreated or treated wild-type and ΔTbPat bloodstream parasites. n = 100 lipid droplets/condition. **c,** Quantification of the mean number of lipid droplet per parasite in bloodstream wild-type and ΔTbPat bloodstream parasites without or with oleate treatment. n = 100 parasites/condition. **d,** Quantification of triacylglycerols (left), cholesteryl esters (middle) and diacylglycerols (right) abundances in wild-type and ΔTbPat bloodstream parasites with or without oleate treatment. Abundances were normalized to 10^8^ cells. n = 3 biological replicates. **e,** Growth curves of wild-type and ΔTbPat bloodstream cells without or with oleate treatment. The curves represent the mean growth of 3 biological replicates. **f,** Volcano-plot of differentially expressed proteins in ΔTbPat bloodstream parasites compared to wild-type cells**. g,** Top upregulated Biological Processes GO terms in ΔTbPat bloodstream cells. **h,** Schematic representation of Log2 fold change of metabolic proteins expression in ΔTbPat parasites. Dashed lines indicate transport processes. Grey circles indicate no significant fold change in protein expression. 1, hexokinase; 2 glucose-6-phosphate isomerase; 3, phosphofructokinase; 4, aldolase; 5, triosephosphate isomerase; 6, glycerol-3-phosphate dehydrogenase; 7, glycerol kinase; 8, glyceraldehyde 3-phosphate dehydrogenase; 9, phosphoglycerate kinase; 10, phosphoglycerate mutase; 11, enolase; 12, pyruvate kinase 1; 13, pyruvate phosphate dikinase; 14, Phosphoenolpyruvate carboxykinase; 15, glycosomal malate dehydrogenase; 16, glycosomal fumarate hydratase; 17, glycosomal NADH-dependent fumarate reductase; 18, pyruvate dehydrogenase E1 α subunit; 19, pyruvate dehydrogenase E1 β subunit; 20, dihydrolipoamide acetyltransferase; 21, pyruvate dehydrogenase complex E3; 22, succinyl-CoA:3-ketoacid coenzyme A transferase; 23, acetyl-CoA hydrolase; 24, succinyl-CoA synthetase α; 25, succinyl-CoA ligase β; 26, mitochondrial NADH-dependent fumarate reductase; 27, mitochondrial fumarate hydratase; 28, mitochondrial malate dehydrogenase; 29, citrate synthase; 30, aconitase; 31, isocitrate dehydrogenase; 32, 2-oxoglutarate dehydrogenase E1 component; 33, 2-oxoglutarate dehydrogenase E1 component; 34, 2-oxoglutarate dehydrogenase E2 component; 35, L-proline dehydrogenase; 36, pyrroline-5-carboxylate dehydrogenase; 37, glutamate dehydrogenase; 38, FAD-dependent glycerol-3-phosphate dehydrogenase; 39, NADH dehydrogenase; 40, Alternative oxidase; 41, ATP synthase F1 subunit α subunit; 42, ATP synthase F1 subunit β subunit; 43, ATP synthase F1 subunit γ subunit; 44, L-threonine 3-dehydrogenase; 45, 2-amino-3-ketobutyrate coenzyme A ligase; 46, Acetyl-CoA synthetase; 47, Acetyl-CoA carboxylase; 48, Elongases 1 – 4. **i**, Schematic representation of the experimental setup for wild-type and ΔTbPat differentiation from slender bloodstream form to procyclic from and subsequent behavior of each cell line. **j**, growth curve of stumpy BSF WT and ΔTbPat cells in glucose-free SDM79 supplemented with 6 mM Cis-aconitate. **k**, Western-blot analysis of PAD1 and EP of WT and ΔTbPat BSF slender form, high-density BSF stumpy form and cells after 96 hours in differentiation medium. Enolase was used as a loading control. OA, oleic acid. Scale bars: 5 µm.

To investigate the underlying adaptive mechanism observed in the lipidomic data, we performed proteomic analyses on WT and ΔTbPat cells in both PCF and BSF stages. In PCF trypanosomes, loss of TbPat caused only minor changes in the proteome with no particular compartment or biological process affected (Extended Data Fig. 4a, Supplementary Table 1). In contrast, BSF parasites showed a much stronger response to TbPat inactivation, suggesting higher sensitivity in this life stage. Strikingly, ΔTbPat BSF cells grown in standard conditions exhibited a global upregulation of mitochondrial enzymes, especially those involved in the TCA cycle and acetate metabolism — enzymes typically more abundant in PCF cells due to their more active mitochondrial metabolism (Fig. 5f, Fig. 5g, Fig. 5h). To assess whether these changes translated into altered metabolic output, we performed quantitative ^1^H-NMR analysis of excreted metabolites from metabolism of the main carbon sources (proline, glucose and threonine) used by trypanosomes. However, no significant differences were found between WT and ΔTbPat BSF cells (Extended Data Fig. 4b). Furthermore, we did not detect any upregulation of electron transport chain components, which remain inactive in BSF forms while being essential for ATP production in PCF cells^44^, indicating that oxidative phosphorylation was still not functional.*T. brucei* defined by an upregulated mitochondrial metabolic proteome could correspond to a population recently been described in parasites isolated from cattle^45^.

The activation of the mitochondrion in bloodstream trypanosomes is a key feature of the differentiation process towards procyclic forms. We hypothesized that in ΔTbPat BSF, the remodeling of the proteome driven by the inability to mobilize lipids could be translated into effects on the transformation from BSF to PCF. To test this, we attempted to differentiate our mutant into PCF as previously described^46^ (Fig. 5k). As shown, the WT pleomorphic cell-line is fully able to express the non-dividing stumpy form marker PAD1 and the PCF marker EP^46^ and grow in the differentiation medium (Fig. 5k), showing its capabilities to differentiate into the insect stage (Fig. 5i). This could not be observed with the ΔTbPat BSF cell line, where parasites consistently die in the differentiation medium and exhibit weaker PAD1 and EP expressions, although the direct cause for this inability to differentiate remains to be investigated (Fig. 5h, Fig. 5k).

In summary, our data point to a strong link between neutral lipid availability and mitochondrial remodeling mediated through TbPat TAG lipase activity, a connection also recently highlighted in yeast and mammalian cells^47^. We show for the first time the essential role of lipid droplets in kinetoplastid lifecycle completion.

## Discussion

This study presents the first functional characterization of a unique orthologue of the lipolysis-limiting enzyme ATGL in the protozoan parasite *Trypanosoma brucei*, which localizes to lipid droplets and modulates parasite metabolism homeostasis and differentiation.

Lipid droplets are conserved key organelles for lipid storage, buffering metabolic stress, supporting cell adaptation to environmental changes and many other functions. Despite their importance in eukaryotic cell biology, knowledge about their dynamics and associated proteins in trypanosomatids remains limited. Here, we demonstrated that TbPat localizes to the LD surface and becomes upregulated under lipid-rich conditions, a phenotype never observed before in trypanosomes. The apparent evolutionary conservation allows parallels between protozoan and metazoan cells and points to conserved LD regulation mechanisms that could be investigated at the single-autonomous-living-cell level. Trypanosomes are attractive model to study biological processes such as RNA editing, glycophosphatidylinositol anchoring, trans-splicing and antigenic variation, all biological phenomena that were initially discovered in these parasites^22^. We prove here that *T. brucei*, as an ancestral living-cell, is a suitable new model to study LD biology.

The existence of an environmental-regulated fatty acid mobilization pathway has already been partially elucidated in trypanosomes. Indeed, previous work has shown that in lipid-poor conditions, the entire fatty acid elongase (ELO) pathway is upregulated to enhance fatty acid synthesis^25^ and the acetyl-CoA carbolyxase (TbACC) enzyme is downregulated in a lipid-rich environment through phosphorylation^48^. While TbPat appears to be regulated in the opposite direction, all these enzymes may be part of a yet uncovered shared mechanism carefully controlling fatty acid homeostasis in cells.

Although TbPat is a functional TAG lipase, its inactivation did not result in a growth defect under standard culture conditions. This suggests that lipid droplet catabolism is not essential under nutrient-rich conditions, but may become critical under stress or in lipid-restricted environments. It is therefore plausible that TbPat and more generally lipid droplet metabolism may become essential *in vivo*, where access to host-derived lipids is limited or tightly regulated. In fact, while TbPat had the same localization and activity across different life stages of *T. brucei*, its inactivation in mammalian stage parasites had much more dramatic effects at the proteomic and lipidomic levels, where cells exhibited a mitochondrial metabolic proteome expression and neutral lipids distribution that resembled more a procyclic-like phenotype. In the mammalian host, *T. brucei* colonizes both the blood and interstitial spaces in tissues; notably, the adipose tissue is a major reservoir for the parasite^49^. In this life stage, *T. brucei* adapted to lipid-rich environments and mostly relies on a continuous supply of lipids to feed its own pathways, likely with the involvement of lipid droplets and TbPat for storage and regulation of the distribution of fatty acids in the cell. Disruption of this careful balance between storage and utilization of lipids may cause a more important stress in mammalian forms than insect stage parasites, which can adapt their metabolism depending on available substrates and can synthesize most of their own lipids in poor environments. TbPat appears central to this balance, as disruption of the neutral lipid mobilization by its inactivation results in the inability of bloodstream parasites to complete their lifecycle since *T. brucei* is unable to differentiate into insect-stage forms. TbPat and more generally lipid droplets thus appear essential in cellular differentiation.

Interestingly, our microscopy data revealed that TbPat-positive LD frequently clustered near the mitochondrion and glycosomes of *T. brucei* (Extended Data Fig. 1c) suggesting potential cross-talk between these organelles. LD-mitochondrion and LD-peroxisome contact sites have been implicated in fatty acid trafficking and β-oxidation in other organisms^39^. While *T. brucei* lacks classical β-oxidation pathways, FFA released from LDs may still support mitochondrial function or glycosomal metabolism by fueling acetate production or other intermediates. Classical and super-resolution microscopy also revealed an uneven distribution of endogenous TbPat at the LD surface, a phenotype which, to our knowledge, has not been observed before in LD-associated lipases. This observation suggests a specific interactor of TbPat either on LD or in contact with other compartments, regulating its spatial distribution and potentially its abundance on the LD monolayer. As the lipolysis rate-limiting enzyme in humans, ATGL is under strict regulation by other proteins to control its activity and maintain cellular homeostasis. A major regulator of ATGL is its binding partner CGI-58/ABHD5, an αβ-hydrolase that activates and significantly increases ATGL lipolytic activity^50^. This interaction is itself regulated by members of the Perilipin family, major LD-associated proteins responsible for protecting the organelle from lipolysis^51^. Identifying and characterizing potential orthologs of these proteins in *T. brucei* must be the focus of further studies to better understand TbPat regulation at the protein level and more generally LD dynamics in trypanosomes.

Based on our results, we propose a model for neutral lipid and fatty acid availability in trypanosomes through an environmental regulation of TbPat expression (Fig. 6), where changes in the lipid environment may directly or indirectly impact both *TbPat* mRNA and protein stabilities. Other proteins, unidentified yet, may be involved in the regulation of the protein stability and/or catalytic activity of TbPat, modulating lipolysis in *T. brucei* and allowing distribution of fatty acids in the parasite to respond to different cellular processes, including metabolism, tissue/host adaptation, organelle remodeling and differentiation. Upstream and downstream effectors participating in the regulation of this pathway now need be characterized to uncover the many roles of lipid droplets in *T. brucei*’s biology.

**Fig. 6:**
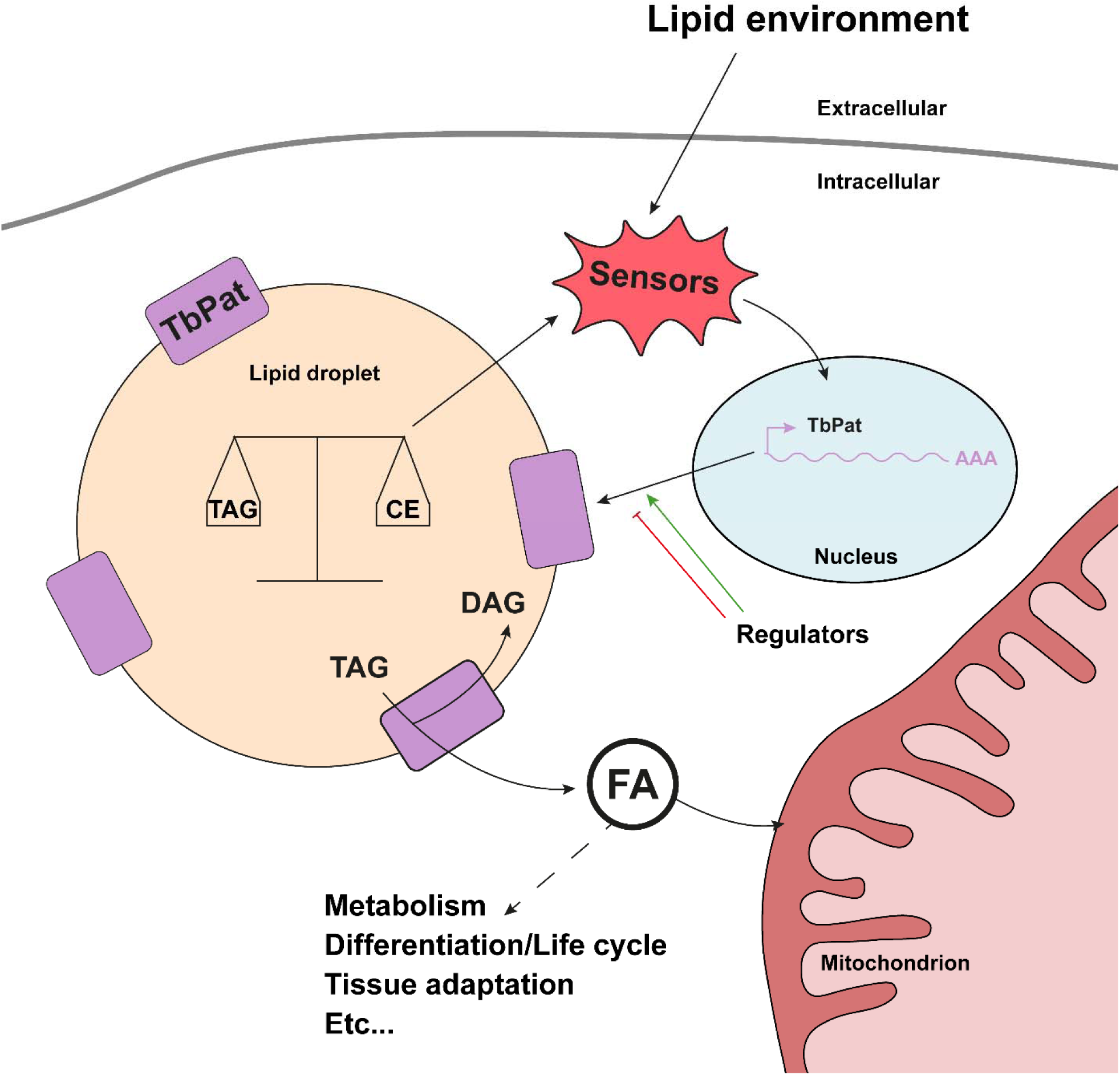
A model for lipid metabolism modulation of *Trypanosoma brucei* biology orchestrated by TbPat. Environmental lipids and the internal lipid droplet neutral lipid balance between triacylglycerol and cholesteryl esters act as sensors to drive the translation of TbPat. Fatty acid synthesis from TAG hydrolysis by TbPat allows maintenance of the cell lipid homeostasis and mitochondrial functions, likely participating in *T. brucei* metabolism, life cycle completion, tissue adaptation, and possibly other biological processes uncovered yet.

In conclusion, we have identified the first functional ortholog of ATGL in protozoan parasites, a patatin-like phospholipase essential for neutral lipid modulation and differentiation in *Trypanosoma brucei*. This work paves the way for the characterization of new lipid droplet-associated proteins and brings *T. brucei* to the forefront as a suitable model to study lipid droplet biology.

## Methods

### Trypanosomes culture and transfections

The procyclic form of *T. brucei* EATRO1125.T7T (TetR-HYG-T7RNAPOL-NEO) was cultured at 27°C with 5% CO_2_ in SDM79 (Semi-Defined Medium)^52^ containing 10% (v/v) heat-inactivated fetal calf serum, 5 μg/ml hemin, 25 μg/ml hygromycin and 10 μg/ml neomycin. The bloodstream form of *T. brucei* AnTat 1.1 90:13 (TetR-HYG-T7RNAPOL-NEO) was cultured at 37°C with 5% CO2 in IMDM (Iscove’s Modified Dulbecco Medium) supplemented with 10 % (v/v) heat-inactivated fetal calf serum, 0.2 mM β-mercaptoethanol, 36 mM NaHCO_3_, 1 mM hypoxanthine, 0.16 mM thymidine, 1 mM sodium pyruvate, 0.05 mM bathocuproine, 1.5 mM L-cysteine, 5 µg/ml hygromycin and 2,5 µg/mL neomycin. Cell growth and fluorescence were monitored daily by cell counting using the cytometer Guava® easyCyte™. All growth curves were made in three independent replicates; graphical representation show the mean of the three replicates. Transfections were performed using the program X-001 of the Amaxa Nucleofector^®^ II and parasites were selected in the appropriate medium.

### Endogenous tagging and inactivation of TbPat with CRISPR/Cas9

Mutations of the endogenous sequence of TbPat were achieved using the CRISPR/Cas9 technology according to a previous paper from our group^33^. For TbPat inactivation, a cassette encoding for the HpaI restriction site surrounded by successive STOP codons and flanked by around 40 bp homologous to the 5’ and 3’ of TbPat surrounding the Cas9 cutting site was used as a donor sequence. A similar approach was used for the endogenous tagging, using a 10TY cassette flanked by the 5’ and 3’ sequences homologous to TbPat. 1.10^6^ parasites were transfected with 1 μg of purified cassette, 30 µg of Cas9 (IDT) and pre-annealing tracrRNA (0.4 µmol) (IDT) and crRNA (0.4 µmol) (IDT). Cells were cloned using a cell sorter (TBMCore Facility, Université de Bordeaux) and selection of double mutants was done by DNA extraction of clonal populations using the prepGEM® Bacteria kit (PBA0500, MicroGEM) followed by PCR amplification. For selection of ΔTbPat homozygotes mutants, the PCR product was digested with HpaI. gRNA were designed using EuPaGDT (http://grna.ctegd.uga.edu/) and TriTrypDB (tritrypdb.org). Primers and crRNA were synthetized by IDT and are listed in Supplementary Table 2.

### Plasmid transfection and inducible expression of TbPat

All plasmid constructs were made in the pLew100 tetracycline-inducible expression vector (phleomycin resistance), into which a GFP was previously inserted (pLew100GFP-X). The TbPat gene was amplified and cloned into NdeI and XbaI (GFP-TbPat and GFP-TbPatΔHD) restriction sites of pLew100GFP-X. Parasites were transfected and selected in SDM79 supplemented with 5 µg/mL of phleomycin. Expression of the fusion protein was induced by adding 1 µg/mL of tetracycline. Primers used for the constructions are listed in Supplementary Table 2.

### BSA-Oleate solution

Stock solutions of 10 mM oleate were prepared by mixing 10 mL of PBS containing 20% of BSA with 32.15 µl of oleate (ThermoFisher 031997.06) drop by drop and shaking at 37°C for 1 hour. The solution was filtered (0.22 µm) and kept at 4°C for short-term storage. For all experiments, the BSA-Oleate solution was added to reach a final concentration of 400 μM in the culture medium.

### Immunofluorescence

Cultured cells were collected by centrifugation and washed with PBS before fixation with 2% paraformaldehyde (PFA) for 10 minutes followed by 0.1 mM glycine quenching for 10 minutes. Cells were spread on slides and permeabilized with 0.05% Triton X-100, followed by a 20 minute-incubation with PBS containing 4% bovine serum albumin (BSA). Cells were incubated for 1 hour with primary antibodies at the indicated dilutions in PBS-BSA 4%: anti-Ty1^34^ 1:250, anti-GFP (Abcam, ab290, 1:2,000 or Abcam, ab13970, 1:500), anti-AceCS^23^ 1:500. Cells were washed with PBS and incubated with secondary antibodies diluted in PBS-BSA 4%: anti-mouse Alexa Fluor 488 (ThermoFisher, A-11001, 1:400), anti-rabbit Alexa Fluor 594 (ThermoFisher, A-11012, 1:400), anti-chicken Alexa Fluor 488 (ThermoFisher, A-11039, 1:400). Slides were washed with PBS and mounted with SlowFade Gold (Molecular Probes). For lipid droplet staining, 0.1 nmol/mL of Lipi-Blue (Dojindo LD-01) was used in colocalization assays and in GFP-tagged strains and 1 μg/mL of Nile Red was used in other experiments. Images were acquired with MetaMorph software on a Leica DM5500 B microscope and processed with ImageJ.

### Cryo-expansion microscopy

U-ExM and Cryo-ExM were performed according to the published protocol^38^. Briefly, for Cryo-ExM, 4.10^6^ cells were loaded on 12 mm coverslips precoated with poly-L-lysine and plunged in liquid ethane. Coverslips were transferred to 5 mL Eppendorf tubes containing 1 mL of acetone, 0.1% PFA and 0.02% glutaraldehyde pre-cooled in liquid nitrogen and placed on dry ice overnight to allow gradual rise of temperature. Cells were then rehydrated with sequential baths of ethanol mixed with 0.1% PFA and 0.02% glutaraldehyde with increasing percentage of water. Cells were embedded following conventional U-ExM protocol^35^. Expanded cells were stained with primary antibodies in PBS-BSA 2% at the following dilutions: anti-Ty1 1:100, anti-BiP^53^ 1:1,000 or anti-GFP 1:500 for 3h at 37°C with agitation. Gels were washed with PBS-Tween 0.1% before incubation with secondary antibodies in PBS-BSA 2%: anti-mouse Alexa Fluor 488 1:400, anti-rabbit Alexa Fluor 594 1:400 or anti-rabbit Alexa Fluor 488 (ThermoFisher, A-11012, 1:400) for 3h at 37°C and washed. NHS-Ester staining (Sigma-Aldrich, 08741, 2 μg/mL) was performed in PBS for 1.5 hours at room temperature. Gels were washed with PBS before re-expansion with water. Images were acquired using a Leica DMI6000 TCS SP8 X confocal microscope and deconvoluted using Huygens Professional 17 with 25 iterations.

### Site-directed mutagenesis

The serine 68 of TbPat was mutated to an alanine using the pLew100GFP-TbPat plasmid as a template. Two complementary primers were synthetized by IDT (Table S1) and amplification was performed using the *Pfu* ultra polymerase (Aligent, 600380). The PCR product was digested with DpnI and transformed in *E. coli* XL1-Blue competent cells. Plasmids were then extracted and sequenced (Eurofins Genomics) to validate the presence of the mutation.

### Reverse transcription

Total RNA was extracted from 2 × 10□ cells using 1 mL of TRIzol reagent (Life Technologies), and its integrity was assessed by agarose gel electrophoresis. RNA concentration was determined with a NanoDrop spectrophotometer. Five micrograms of RNA were treated with DNase I using the TURBO DNA-free Kit (Ambion), and cDNA synthesis was performed using oligo(dT)_₁₂₋₁₈_ primers (Invitrogen) and SuperScript IV Reverse Transcriptase. The primers used for RT-PCR are listed in Supplemental Table 2.

### Western-blot

Total protein extracts from 5.10^6^ parasites were separated by SDS-PAGE (10% Mini PROTEAN TGX stain-free precast gradient gels, Bio-Rad) and blotted on TransBlot Turbo Midi-size PVDF Membranes (Bio-Rad). Membranes were blocked with PBS-0.05% Tween20 containing 5% of skimmed milk powder. Primary and secondary antibodies were diluted in PBS-Tween-Milk as followed: anti-Ty1 1:500, anti-Enolase^54^ 1:100,000, anti-GFP 1:1,000, anti-PFR^55^ 1:10,000, anti-EP^46^ 1:400, anti-PAD1 1:1,000. Secondary antibodies conjugated to horseradish peroxidase (anti-mouse HRP, Bio-Rad, 1706516, 1:5,000, anti-rabbit HRP, Bio-Rad, 1706515, 1:10,000) allowed revelation using the Clarity Western Enhanced-Chemiluminescence Subtrate (Bio-Rad) according to the manufacturer’s instructions. Images were acquired with the ImageQuant Las 4000 imager (GE Healthcare) and processed with ImageJ.

### Lipidomics analysis

Parasites (at least 5.10^7^ cells per replicate, three replicates per condition) were metabolically quenched by rapid chilling in a dry ice-ethanol slurry bath and centrifuged at 4°C. Pellets were washed three times with ice-cold PBS and transferred to a microcentrifuge tube for storage at –80°C until analysis. Total lipid samples were spiked with 20 nmol C21:0 phosphatidylcholine and extracted by chloroform:methanol, 1:2(v/v) and chloroform:methanol, 2:1 (v/v). The pooled organic phase was subjected to biphasic separation by adding 0.1% KCl and was then dried under N2 gas flux prior to being dissolved in 1-butanol. For the total fatty acid analysis, an aliquot of the lipid extract was derivatized on-line using MethPrep II (Alltech) and the resulting FA methyl esters were analyzed by GC-MS as previously described (26). For the quantification of each lipid, total lipid was separated by 2D HPTLC using chloroform/methanol/28% NH4OH, 60:35:8 (v/v/v) as the 1st dimension solvent system and chloroform/acetone/methanol/acetic acid/water, 50:20:10:13:5 (v/v/v/v/v) as the 2nd dimension solvent system. Each lipid spot was extracted for quantification of fatty acids by gas chromatography-mass spectrometry (Agilent 5977A-7890B) after methanolysis. Fatty acid methyl esters were identified by their mass spectrum and retention time and quantified by Mass Hunter Quantification Software (Agilent) and the calibration curve generated with fatty acid methyl esters standards mix (Sigma CRM47885). Each lipid content was normalized according to parasite cell number and a C21:0 internal standard (Avanti Polar lipids). Lipids were analyzed by LCMSMS as previously described^56^. Briefly, dried down samples were reconstituted in 80 μL methanol and incubated at 30°C for 5 minutes with vigorous vortex. Samples (1 μL) were analyzed by LCMS (Agilent 1290 infinity/Infinity II Agilent) MS (Agilent 6495c triple quadrupole) in our platform GEMELI. Acquisition DMRM (dynamic multiple reaction monitoring) method used was referred to as described previously^57^ with a modification for *Trypanosoma* lipid species. The resulting LCMS data was subjected to targeted analysis using Mass Hunter Quantification software (Agilent). Each lipid species was quantified using a calibration curve of each representative lipid with known abundance. Then each lipid abundance was normalized according to the cell ratio.

### Mass spectrometry and data analysis

1.10^8^ parasites/replicate were washed in their culture medium without serum. Pellets were then resuspended in 200 µL of a lysis solution (4% of sodium deoxycholate in 100 mM Tris-HCl pH 8 and a protease inhibitor cocktail) prepared extemporaneously and cooled to 4°C. Solutions were warmed to 95°C for 5 minutes and sonicated at 4°C for 5 cycles of 30 seconds at maximum output. Protein samples were solubilized in Laemmli buffer and 5 µg of proteins were deposited onto an ultra-short preparative SDS-PAGE gel. A single band, containing the complete sample proteome, was excised and subjected to overnight trypsin digestion. The resulting peptides were extracted, concentrated, and analyzed by LC-MS/MS using an Ultimate 3000 nanoLC system (Dionex, Amsterdam, The Netherlands) coupled to an Electrospray Orbitrap Fusion™ Lumos™ Tribrid™ Mass Spectrometer (Thermo Fisher Scientific, San Jose, CA). Data acquisition was performed in data-independent acquisition (DIA) mode. Each duty cycle consisted of one full scan (resolution: 60,000; AGC target: 4E5; maximum injection time: 50 ms; mass range: 350−1400 m/z), followed by 35 DIA MS/MS scans (resolution: 30,000; AGC target: 5E5; maximum injection time: 54 ms) covering the mass range of 380−980 m/z with variable-width isolation windows. HCD collision energy was set to 25%.

DIA data were processed using Chimerys within Proteome Discoverer 3.1 (Thermo Fisher Scientific Inc.) and searched against the *Trypanosoma brucei* protein database (strain 927 – 9,243 entries). Only high-confidence peptides, corresponding to a 1% false discovery rate (FDR) at the peptide level, were retained. Peptide quantification was performed using the Fragment Ions Quantifier node, with normalization based on the total *Trypanosoma* peptide amount. Protein ratios were calculated as the median of all possible pairwise peptide ratios. A t-test was applied based on the background population of peptides or proteins and adjusted using the Benjamini-Hochberg correction. Quantitative data were considered only for proteins identified with a minimum of two peptides and a statistical p-value below 0.05. To analyze for differential abundance of proteins in the conditions, proteins were filtered based on several criteria: i) the protein is a master protein, ii) it has more than one unique peptide, iii) it is present in all replicates of both conditions or completely absent in one of the conditions and iv) the mean abundance of the protein is superior to 5.10^5^ in at least one condition. Proteins significantly downregulated or upregulated in ΔTbPat PCF or BSF cells and their description/Curated GO Terms are summarized in Extended Data Table 1. Top Biological Processes GO Terms for the upregulated proteins in ΔTbPat BSF cells were sorted by a minimum cutoff of 3 proteins per GO terms. All GO Terms can be found in the Extended Data Table 1. Graphical representations were made using R version 4.4.1 with the ggplot2 and ggrepel packages.

### NMR spectrometry

2×10^7^ *T. brucei* BSF cells were collected by centrifugation at 1,400 x g for 10 min, washed twice with PBS supplemented with 2 g/L NaHCO_3_ (pH 7.4) and incubated in 1.5 mL (single point analysis) of PBS-NaHCO_3_. Cells were incubated in a buffer containing one ^13^C-enriched carbon source (4 mM, U-^13^C-Glucose alone or with U-^12^C-Proline or U-^12^C-Threonine) for 90 minutes. The integrity of the cells during the incubation was checked by microscopic observation. The supernatant (1.5 mL) was collected and 50 μL of maleate solution in Deuterated water (D_2_O, 10 mM) was added as an internal reference. ^1^H-NMR spectra were performed at 500.19 MHz on a Bruker Avance III 500 HD spectrometer equipped with a 5 mm cryoprobe Prodigy. Measurements were recorded at 25°C. Acquisition conditions were as follows: 90° flip angle, 5,000 Hz spectral width, 32 K memory size, and 9.3 sec total recycle time. Measurements were performed with 64 scans for a total time close to 10 min 30 sec. Results are available in Extended Data Table 3 and representative data in Extended Data Fig. 4.

### *In vitro* differentiation of bloodstream parasites

Dividing slender parasites were grown in IMDM medium for 2 days to reach high-density (more than 1.10^6^ cells/ml) and allow differentiation into the non-dividing stumpy form. Parasites were then centrifuged and 5.10^5^ cells were resuspended into glucose-free SDM79 medium^58^ supplemented with 6 mM cis-aconitate (differentiation medium) (Sigma, A3412). Cells were maintained at 27°C and growth was assessed every day. Parasites were taken at different time points for anti-PAD1 and anti-EP Western-blot analysis: before high-density culture, 2 days of high-density culture and 4 days after incubation in the differentiation medium. The experiment was performed in 3 independent replicates.

### Statistical analysis

All statistical analyses were performed using Prism (GraphPad) software v8.0.2. Results are presented as mean ± standard deviation. To get the mean number of lipid droplets per parasite, 100 parasites/condition were manually analyzed. To quantify the lipid droplet area, 100 droplets were analyzed for each condition. Results were subjected to either Student’s t-test (comparison of two groups) or one-way/two-way ANOVA (comparison of four groups) with Tukey’s multiple comparison test to determine statistical differences between groups. p-value (confidence interval of 95%) is indicated as ns (not significant), * (<0.05), ** (<0.01), *** (<0.001) or **** (<0.0001).

## Acknowledgments

Image acquisition on the confocal microscope was performed in the Bordeaux Imaging Center (BIC), a service unit of the CNRS-INSERM and Université de Bordeaux, member of the national infrastructure France BioImaging supported by the French National Research Agency (ANR-10-INBS-04). We thank Mónica Fernández Monreal from the BIC for access to the plunger and ethane for cryo-fixation of trypanosomes. Cell sorter analyses were performed at the TBMCore facility (FACSility) on BD FACSMelody™. We thank Harald Wodrich and Nicolas Landrein from the MFP SpacVir team for giving us access to their confocal microscope. Finally, we thank Bénédicte Salin and Corinne Blancard from the IBGC CNRS research unit for the many electron microscopy trials.

## Funding

The iMET team (LR and/or FB) and CYB are funded by the Centre National de la Recherche Scientifique (CNRS, https://www.cnrs.fr/), the Agence Nationale de la Recherche (ANR, https://anr.fr/) through the ParaFrap “Laboratoire d’Excellence” (LabEx, https://www.enseignementsup-recherche.gouv.fr/cid51355/laboratoires-d-excellence.html) (ANR-11-LABX-0024, attributed to French parasitologists including FB and CYB), and ANR OIL (ANR24-CE15-2171-01 attributed to LR and CYB). Proteomic analysis was supported through a grant from the Département des Sciences Biologiques et Médicales of the Université de Bordeaux “AAP SBM 2024 Utilisation d’une plateforme ou plateau technique” attributed to PH. In addition, the iMET team is supported by the Université de Bordeaux (https://www.u-bordeaux.fr/), the “Fondation pour la Recherche Médicale” (FRM, https://www.frm.org/) (“Equipe FRM,” grant no. EQU201903007845), ANR ADIPOTRYP (ANR19-CE15-0004-01) attributed to FB. CYB and YYAB were also supported by Agence Nationale de la Recherche, France (Project ApicoLipiAdapt grant ANR-21-CE44-0010; Project Apicolipidtraffic grant ANR-23-CE15-0009-01; Project OIL grant ANR-24-CE15-2171-02), The “Fondation pour la Recherche Médicale” (FRM EQU202103012700), LIA-IRP CNRS Program (Apicolipid project), the Université Grenoble Alpes (IDEX ISP Apicolipid) and Région Auvergne Rhone-Alpes for the lipidomics analyses platform (Grant IRICE Project GEMELI), Collaborative Research Program Grant CEFIPRA (Project 6003-1) by the CEFIPRA (MESRI-DBT).

## Contributions

PH, CYB, ET, FB and LR designed the study. PH and LR performed most of the experiments and analyzed all the data. EB and/or PH performed the expansion microscopy experiments and analyses. KA performed the reserve transcription. YYAB performed the lipidomics analyses at the GEMINI platform (Grenoble). SC performed the proteomic analysis at the Bordeaux Proteome platform. MB performed the ^1^H-NMR analysis and quantifications at the CRMSB (Bordeaux). LR supervised the project. PH and LR wrote the manuscript with input from all the authors.

---

**Extended Data Fig. 1:**
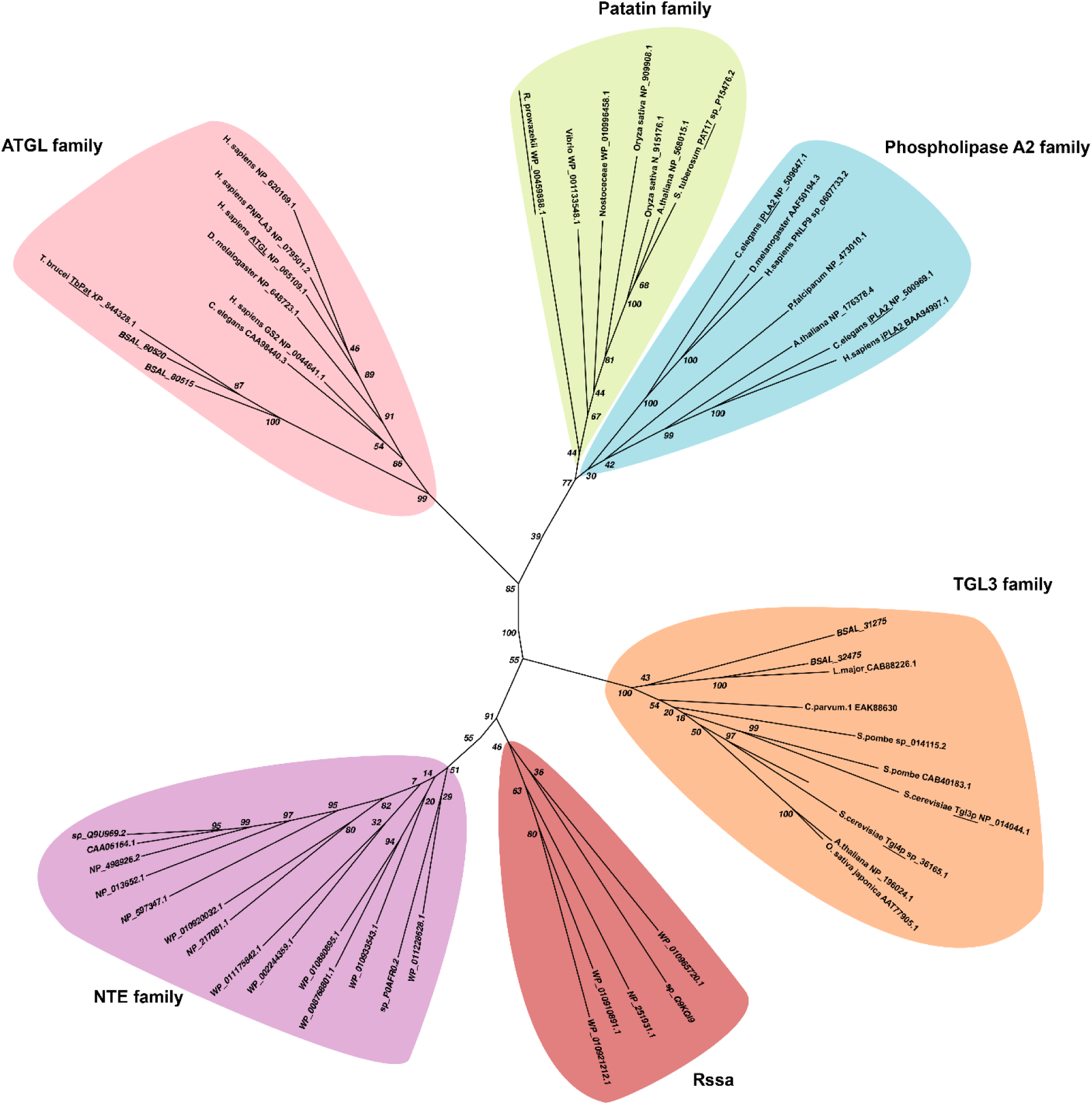
TbPat clusters with Adipose Triglyceride Lipase family. Maximum likelihood phylogeny of Patatin-like proteins. An unrooted tree was built in IQ-TREE by using the best-fit model with 1000 replicates for bootstrapping. Sequences were retrieved from Smirnova & al^7^ and combined with TbPat. Proteins sequences with Genbank identifiers are available in Extended Data File 1. Proteins were aligned using Clustal Omega algorithm. Principal Patatin-like families are colored and main representative proteins are underlined.

**Extended Data Fig. 2:**
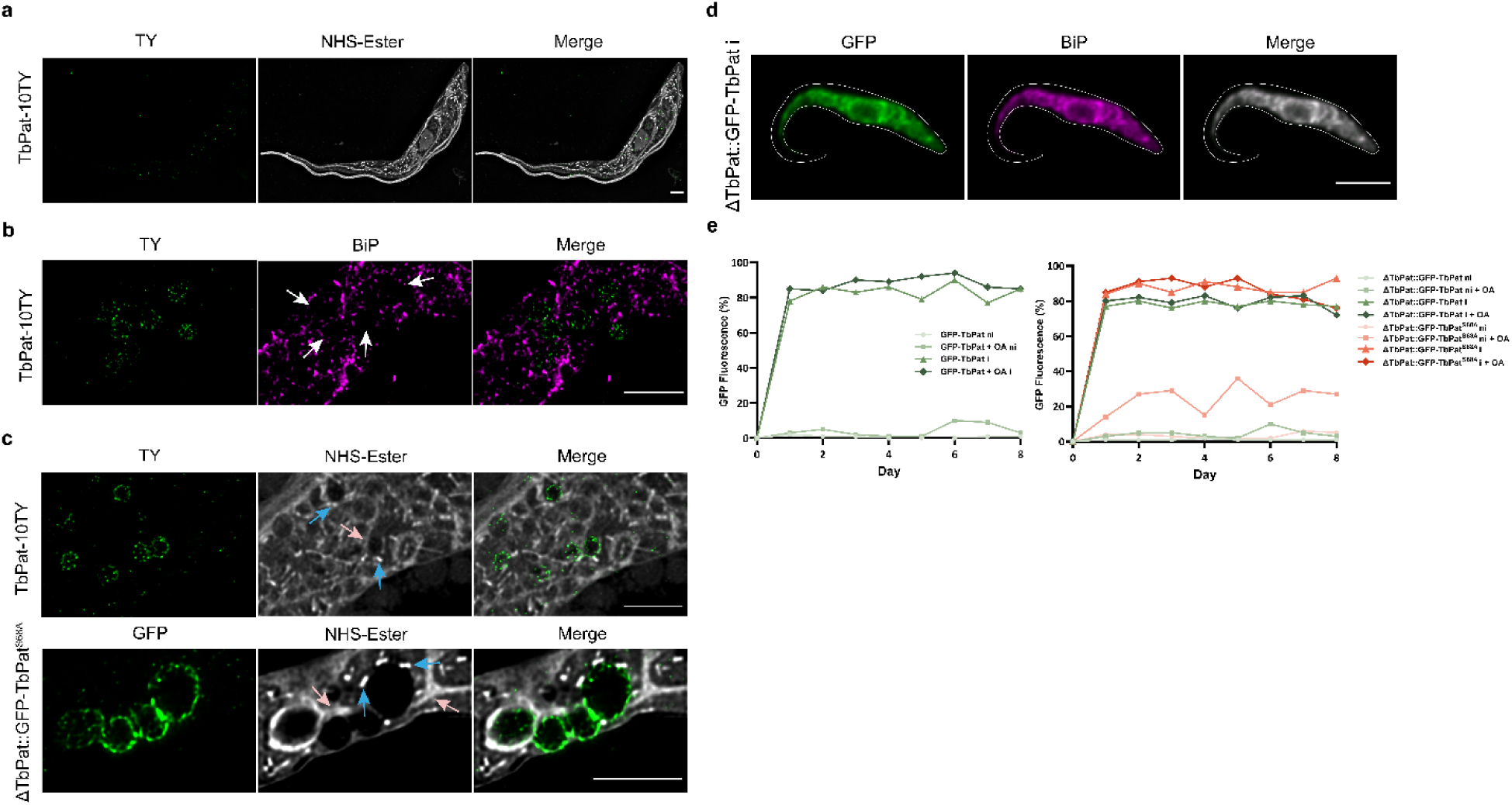
Detailed localization and expression of TbPat-mutant cell lines. **a,** Classical expansion microscopy of TbPat-10TY parasites. **b,** Cryo-expansion microscopy of TbPat-10TY. BiP/HSP70 marks the endoplasmic reticulum. White arrows indicate TbPat-positive lipid droplets. **c,** Cryo-expansion microscopy of TbPat-10TY (top) and ΔTbPat::GFP-TbPat^S68A^ (bottom) induced cells. The blue and pink arrows indicate glycosomes and the mitochondrion, respectively. **d,** Immunofluorescence of ΔTbPat::GFP-TbPat induced cells show colocalization of GFP-TbPat and the endoplasmic reticulum marked by BiP. **e,** Time course expression of GFP fluorescence during the growth curves of wild-type, ΔTbPat, ΔTbPat::GFP-TbPat and ΔTbPat::GFP-TbPat^S68A^ cell lines, monitored daily by flow cytometry. The curves represent the mean fluorescence of 3 biological replicates. Scale bars: 5 µm.

**Extended Data Fig. 3:**
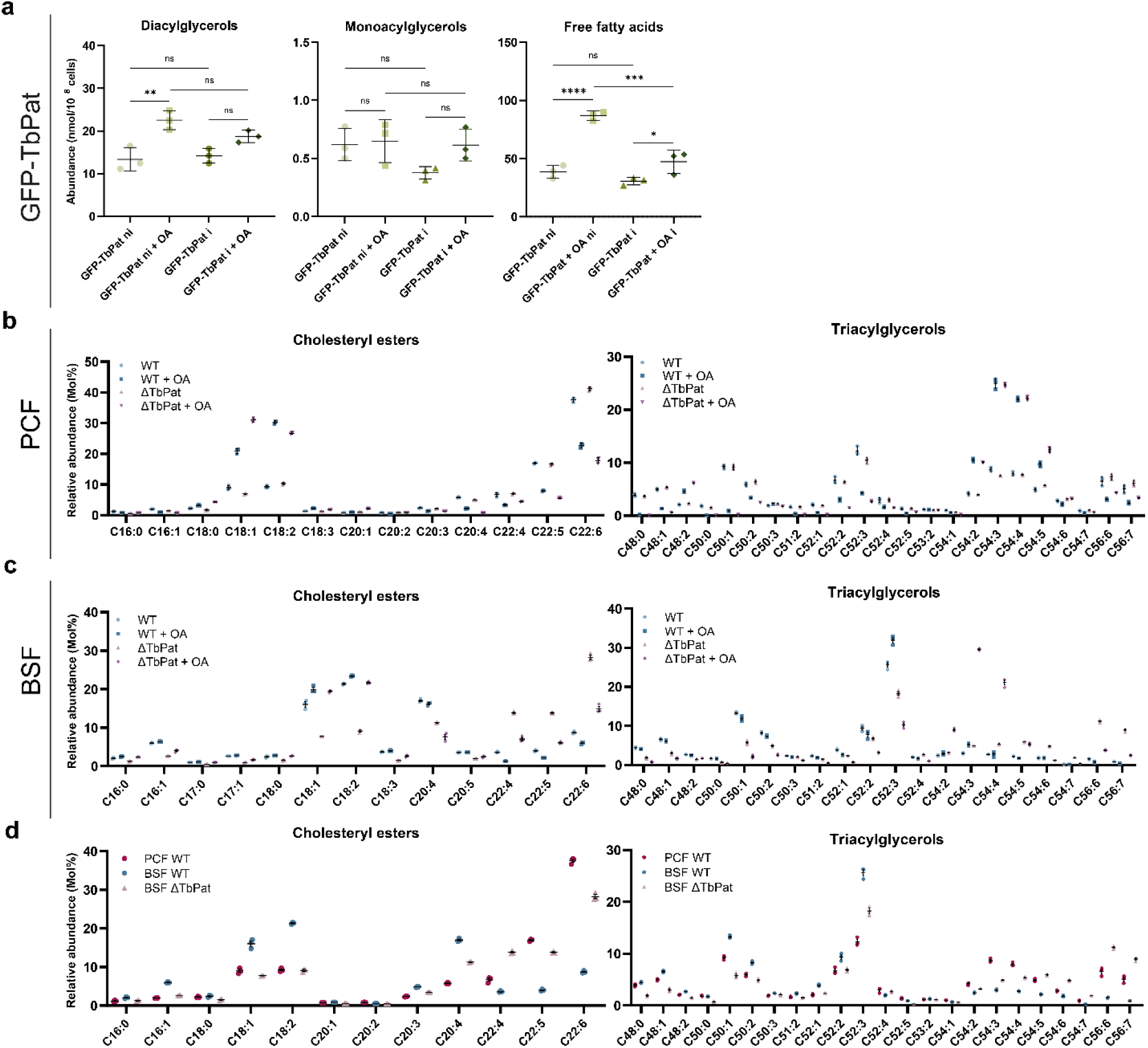
Detailed analysis of lipidomic data from different mutants. **a,** Diacylglycerols, monoacylglycerols and free fatty acids quantifications in non-induced or induced GFP-TbPat without or with oleate treatment for 24 hours. **b, c,** Cholesteryl esters (left) and triacylglycerols (right) molecular species relative abundances profiles in procyclic (**b**) and bloodstream (**c**) wild-type and ΔTbPat cells. **c, d,** Comparison of cholesteryl esters (left) and triacylglycerols (right) molecular species relative abundance profiles between wild-type procyclic and wild-type and ΔTbPat bloodstream cells. For all data, n = 3 biological replicates. ni, non-induced; i, induced; OA, oleic acid, PCF, procyclic; BSF, bloodstream.

**Extended Data Fig. 4:**
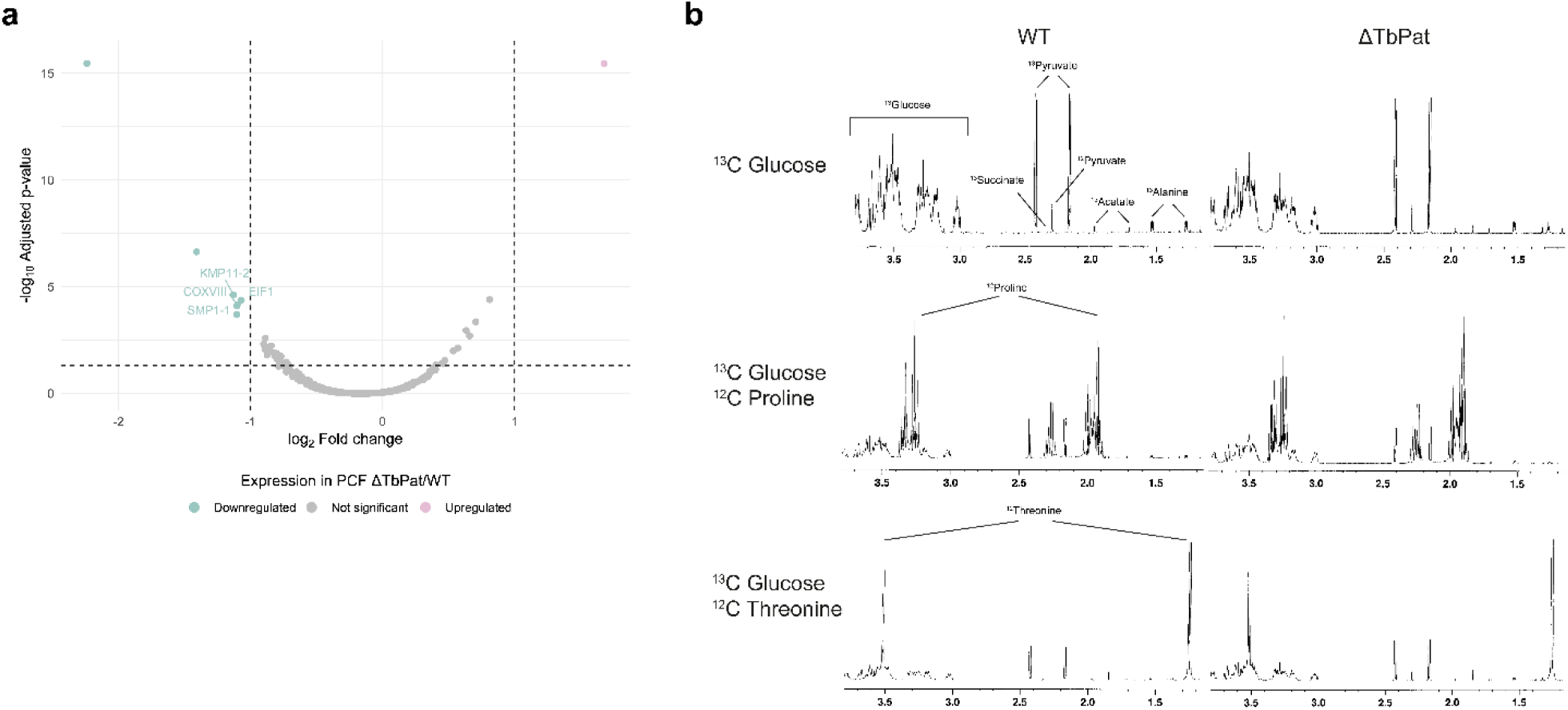
PCF proteomic and BSF Proton (^1^H) NMR analyses. **a,** Volcano-plot of differentially expressed proteins in ΔTbPat procyclic parasites compared to wild-type cells**. b,** Proton (^1^H) NMR analysis and analysis of end products excreted from wild-type and ΔTbPat bloodstream form parasites. 4 mM [U-^13^C]-glucose and 4 mM proline (middle) or 4 mM [U-^13^C]-glucose and 4 mM threonine (bottom). The panel shows a representative NMR spectrum for each condition. Excreted end products are indicated by black lines. Detailed results are available in Extended Data Table 3.

## Notes

### Competing Interest Statement

The authors have declared no competing interest.

